# Self-assembled nanoparticles enable delivery of VCP inhibitor, resulting in enhanced photoreceptor protection in retinal explants

**DOI:** 10.1101/2021.06.11.448060

**Authors:** Merve Sen, MD Al-Amin, Eva Kicková, Amir Sadeghi, Jooseppi Puranen, Arto Urtti, Paolo Caliceti, Stefano Salmaso, Blanca Arango-Gonzalez, Marius Ueffing

## Abstract

Mutations in rhodopsin lead to its misfolding resulting in autosomal dominant retinitis pigmentosa (adRP). Pharmacological inhibition of the ATP-driven chaperone valosin-containing protein (VCP), a molecular checkpoint for protein quality control, slows down retinal degeneration in animal models. However, poor water-solubility of VCP inhibitors poses a challenge to their clinical translation as intravitreal injections for retinal treatment. In order to enable the delivery of VCP inhibitors, we have developed and investigated two formulations for the VCP inhibitor ML240. Nanoformulations of ML240 were obtained by using amphiphilic polymers methoxy-poly (ethylene glycol)_5kDa_-cholane (mPEG_5kDa_-cholane) and methoxy-poly (ethylene glycol)_5kDa_- cholesterol (mPEG_5kDa_-cholesterol). Both formulations increased the water-solubility of ML240 by two orders of magnitude and prolonged the drug released over ten days. Encapsulation of ML240 in mPEG_5kDa_-cholane showed superior photoreceptor protection at lower drug concentrations, normalized rhodopsin localization, and alleviated inflammatory microglial responses in an *ex vivo* rat model of retinal degeneration. The study demonstrates the potential of VCP inhibitor nanoformulations to treat adRP, a pharmacologically orphan disease.

## 2. Introduction

Retinitis Pigmentosa (RP) is a group of hereditary visual disorders characterized by progressive loss of photoreceptor neurons, eventually leading to blindness [1]. More than 3,000 mutations in 70 different genes causing RP have been identified, showing variations in age of onset, rate of progression, and clinical outcomes [2]. Nevertheless, night blindness caused by degeneration of rod photoreceptors is a typical early clinical manifestation of RP, followed by progressive loss of peripheral vision and ultimately loss of central vision due to degeneration of cone photoreceptors [1].

Mutations in rhodopsin (*RHO*) are a common cause of RP [3,4] as at least 150 known point mutations have been previously reported to cause RP [5,6]. A point mutation in the *RHO* N-terminus where proline is replaced by histidine (*RHO*^*P23H*^) is the most common genetic defect in the North American population, resulting in an autosomal dominant form of RP (adRP) [7] with altered folding of rhodopsin that delays with its physiological progress from the endoplasmic reticulum (ER) to the plasma membrane [8,9]. Although the precise mechanism by which RHO^P23H^ triggers RP is not fully understood, it is known that misfolded RHO^P23H^ forms a complex with the ER effector valosin-containing protein (VCP) in mammalian cells [8]. The interaction of RHO^P23H^ with VCP promotes its translocation from the ER and degradation by the proteasome. Accordingly, inhibition of VCP function has been shown to strongly mitigate retinal degeneration in a *Drosophila* model with a similar mutation [10]. We have also shown that chemical inhibition and downregulation of VCP activity rescue degenerating P23H photoreceptors and improves their functions in the RHO^P23H^ rodent model of RP both *in vitro* and *in vivo* [11,12].

Even though inhibition of VCP is a promising therapeutic approach for RP, the poor solubility of known VCP inhibitors in water represents a major obstacle to its clinical translation. Furthermore, the delivery of therapeutics to the retinal targets is hampered by ocular barriers, necessitating intravitreal injections (IVT) required for retinal drug delivery [13]. Intravitreally injected small molecules, such as VCP inhibitors, are rapidly cleared across the blood-ocular barriers to the systemic circulation within less than one day [11,14]. Frequent invasive injections to the vitreal cavity are not easily feasible in clinical practice for safety and logistics reasons. Therefore, efficient delivery systems are needed to overcome the poor biopharmaceutical properties of VCP inhibitors and enable sustained drug release in the vitreous.

Intravitreal nanocarriers have shown prolonged residence in the vitreous in several studies [15-18]. In addition, nanocarriers (e.g., liposomes of 50 nm in diameter) are able to permeate into the retina across the inner limiting membrane [19]. Thus, nanocarriers could be suitable for formulating low soluble VCP inhibitors. A well-designed polymeric drug delivery system could not only increase the solubility of VCP inhibitors but also prolong their residence in the vitreous and sustain drug delivery into the retina.

Monomethoxy-polyethylene glycol (mPEG)-cholesterol is an amphiphilic material that has been exploited in drug delivery systems to formulate a variety of drugs to treat diseases of the central nervous system, infectious diseases, and cancer [20,21]. We have shown that self-assembling polymers obtained by end-functionalization mPEG with cholanic acid, a polycyclic molecule structurally similar to cholesterol, can be successfully used for the delivery of proteins, peptides, and low molecular weight drugs [22-26].

Since mPEG-cholane and mPEG-cholesterol are able to increase the solubility of poorly soluble drugs by forming self-assembled nanosystems, they are promising candidates for the retinal delivery of VCP inhibitors. Therefore, we explored mPEG_5kDa_- cholesterol and mPEG_5kDa_-cholane for intravitreal delivery of VCP inhibitors, potentially increasing drug solubility, prolonging intravitreal residence, and improving their controlled release. One of the most potent VCP inhibitors (ML240) was loaded into mPEG_5kDa_-cholesterol and mPEG_5kDa_-cholane colloidal formulations. Drug solubilization, retinal penetration, and therapeutic efficacy were investigated. We show here that nanoformulation improves the efficacy of ML240 in organotypic cultures of RHO^P23H^ rats, a model known to mimic human adRP.

## 3. Material and Methods

### Study approval and animals

All *in vitro* procedures were approved by the Animal Welfare Commission of the Tuebingen University committee on animal protection (§4 registrations from 26.10.2018 AK 15/18 M) and performed in compliance with the Association for Research in Vision and Ophthalmology ARVO Statement on animal use in ophthalmic and vision research. All efforts were made to minimize the number of animals used and their suffering. Homozygous P23H transgenic rats (produced by Chrysalis DNX Transgenic Sciences, Princeton, NJ) of the line SD-Tg(P23H)1Lav (P23H-1) were kindly provided by Dr. M. M. LaVail (University of California, San Francisco, CA) or the Rat Resource and Research Center (RRRC) at the University of Missouri. Animals were housed in the animal facilities of the Institute for Ophthalmic Research under standard white cyclic lighting, with access to food and water ad libitum. To reflect the genetic background of adRP, we employed heterozygous P23H rats obtained by crossing homozygous P23H transgenic rats with wild-type rats (CDH IGS Rat; Charles River, Germany).

### ML240 encapsulation in mPEG_5kDa_-cholane and mPEG_5kDa_-cholesterol-based particles

mPEG_5kDa_-cholane and mPEG_5kDa_-cholesterol were synthesized according to published protocols [26,27].

The ML240-loaded nanoparticles were generated according to the “film hydration” approach [28]. Briefly, 0.6 mg/mL of ML240 (Tocris Bioscience, Bio-Techne GmbH, 5153, Minnesota, US) in 5 mL acetone was mixed with 5 mL of 0-6 mg/mL mPEG_5kDa_- cholane or mPEG_5kDa_-cholesterol in acetone. The organic solvent was removed under reduced pressure using a rotary evaporator for 30 minutes. Then, the polymer/ML240 films were hydrated with 1.5 mL of 10 mM phosphate, 150 mM NaCl (PBS), pH 5.0, resulting in polymer concentrations of 0-20 mg/mL and 2 mg/mL ML240. The mixtures were vortexed and sonicated for two minutes and stirred overnight. Afterward, the pH was adjusted to pH 7.4 with 0.1 M sodium hydroxide and then centrifuged twice at 14,800 rpm for 3 minutes to remove the non-encapsulated ML240. The concentration of ML240 in the supernatant was assessed by RP-HPLC using a Jasco AS-1555 System (Tokyo, Japan), equipped with 2 PU-2080 plus pumps, a spectrophotometric detector UV-2075 plus set at 259 nm, and a Luna 5 µM C18 column (Phenomenex, CA, US) eluted at a flow rate of 1 mL/min with water (eluent A) and acetonitrile (ACN, eluent B), both containing 0.05% of trifluoroacetic acid (TFA), in a gradient mode: 20 min from 40% to 90% eluent B. The concentration of ML240 was derived from a calibration curve obtained by injecting known concentrations of ML240 in acetone (0-80 µg/mL).

### Size, morphology, and stability of ML240-loaded nanoparticles

Nanoparticle dispersions obtained with 10 mg/mL polymers were diluted to 1 mg/mL with PBS, pH 7.4 at 37 ° C, and analyzed by dynamic light scattering (DLS) using a Malvern Zetasizer Nano S system (Malvern, UK). Size and polydispersity were determined on the basis of the intensity signal.

The stability of samples obtained with 10 mg/mL polymers was assessed by incubating 2 mg/mL ML240-loaded nanoparticle dispersion in PBS, pH 7.4 or in 1:1 porcine vitreous/PBS, pH 7.4, at 37 °C. At scheduled times, the samples were diluted to 1 mg/mL with PBS, pH 7.4, and hydrodynamic size and polydispersity were measured as reported above.

Transmission electron microscopy (TEM) analysis was performed using an FEI Tecnai G2 microscope (Hillsboro, OR, USA). ML240 loaded mPEG_5kDa_-cholane, and ML240 loaded mPEG_5kDa_-cholesterol nanoparticles in water were transferred on a carbon-coated copper grid (400 mesh). The excess volume was removed using filter paper, and the samples were negatively stained with 1% uranyl acetate in distilled water and analyzed.

### Release study from polymeric nanoparticles

*In vitro* release study of ML240-loaded nanoparticles was performed at 37 °C in PBS, pH 7.4. Briefly, 1 mL of ML240-loaded mPEG_5kDa_-cholane or mPEG_5kDa_-cholesterol nanoparticles (10 mg/mL) was transferred into a 3.5 kDa molecular weight cut-off float-a-lyzer (Spectrum Laboratories, CA, US) and dialyzed against 2 L of the same buffer. At scheduled time intervals, 20 µL of the samples inside the dialysis device was withdrawn and analyzed by RP-HPLC as reported above to quantify the residual ML240. The release medium was replaced twice a day with fresh medium.

### Preparation of organotypic culture of RHO^P23H^ rat retinal explants

Retinal explants were cultured as previously described [29,30]. Briefly, postnatal day (PN) 9 RHO^P23H^ rats were sacrificed, eyes were removed, and incubated for 15 minutes at 37 °C in a 12% Proteinase K solution (MP Biomedicals, 0219350490). The enzymatic digestion was stopped by the addition of 20% FBS (Sigma-Aldrich, F7524). First, the sclera was carefully removed, leaving the retinal pigmented epithelium attached to the eyeball. Incisions were made at the edge of the cornea to remove the cornea, lens, and hyaloid body. Then, four equidistant, radial incisions were made in the retina. Tissue was transferred to a 0.4 μm transwell polycarbonate membrane insert (Corning Life Sciences, CLS3412), having the RPE side touching the membrane. Retinae were cultivated in Neurobasal A (Gibco, Thermo Fischer Scientific, US, 21103049) supplemented with 2% B-27 supplement (Gibco, Thermo Fischer Scientific, US, 17504044), 1% N2 supplement (Gibco, Thermo Fischer Scientific, US, 17502048), 1% penicillin solution (Gibco, Thermo Fischer Scientific, US, 15140-122), and 0.4% GlutaMax (Gibco, Thermo Fischer Scientific, US, 35050061). Explants were maintained at 37 °C in a humidified 5% CO_2_ atmosphere. The medium was changed every second day. The PN9 cultures were fixed at DIV6, which corresponds to PN15, the peak of degeneration *in vivo* age-matched mutants for short-term treatment. The PN9 cultures were fixed at DIV21 for long-term treatment. Treatments were given either basolaterally as 5 µM ML240-loaded mPEG_5kDa_-cholane or mPEG_5kDa_-cholesterol nanoparticles mixed in Neurobasal A medium below the explant, in the lower compartment. The medium was changed every second day or apically (to mimic dosing via IVT and, e.g., subretinal injections) as 15 µL drops for 5 µM ML240-loaded mPEG_5kDa_-cholane or mPEG_5kDa_-cholesterol nanoparticles carefully applied directly on top of the explant cultures (applied once on the retina and the medium was changed every second day). Drug-free polymeric nanoparticles at equivalent concentrations were used as negative vehicle controls. Untreated Neurobasal A medium was used as a control.

### Fixation and sectioning of RHO^P23H^ rat retinal explants

Treated rat explants were fixed in 4% PFA (Polysciences, Inc., Warrington, PA, USA) at room temperature for 40 minutes. Retinae were then washed with PBS for 10 minutes and cryoprotected by incubation in graded sucrose solutions (10% for 10 minutes, 20% for 20 minutes, and 30% for 40 minutes). Subsequently, tissues were embedded in Tissue-Tek® O.C.T.™ Compound (Science Services GmbH, Munich, Germany). Vertical sections (14 µm) were obtained on a Leica CM3050S Microtome (Leica Biosystems, Wetzlar, Germany), air-dried at 37 °C for 1 hour, and stored at −20°C until use.

### Staining and imaging of RHO^P23H^ rat retinal explants

Terminal deoxynucleotidyl transferase dUTP nick end labeling (TUNEL) assay [31] was performed on cryosections of retinal explants using an *in situ* cell death detection kit conjugated with fluorescein isothiocyanate (Roche, 11684795910). DAPI (Vectashield Antifade Mounting Medium with DAPI; Vector Laboratories, H-1200) was used as a nuclear counterstain.

For rhodopsin and Iba1 staining, sections were incubated overnight at 4 °C with an anti-rhodopsin mouse (Sigma-Aldrich, MAB5316, 1:300) or anti-Iba1 rabbit antibody (Fujifilm Wako Chemicals, 019-19741, 1:200), respectively. Fluorescence immunocytochemistry was performed using Alexa Fluor™ 568 dye-conjugated goat anti-mouse IgG (Molecular Probes, A-11031, 1:500) or Alexa Fluor™ 488 dye-conjugated goat anti-rabbit IgG (Cell Signaling Technology, A 11034, 1:500). All mounted sections were imaged using a Zeiss Axio Imager Z1 ApoTome microscope equipped with a Zeiss AxioCam digital camera and AxioVision 4.7 software.

### Fundus camera and optical coherence tomography (OCT)

The *in vivo* animal experiments were approved according to the ethical guidelines of the University of Eastern Finland (license number ESAVI-2020-027769). About four months old pigmented male rats (HsdOla:LH) were used for *in vivo* studies. Two rats were used for each formulation. Both eyes of the animal were injected. The imaging was performed before, immediately, and two weeks after injection using OCT and fundus camera (Phoenix MICRON™ MICRON IV/OCT, CA, USA). The injections and imaging were performed under anesthesia using subcutaneous injection of medetomidine (dose of 0.4 mg kg−1, Domitor vet 1 mg/ml; Orion Pharma, Espoo, Finland) and ketamine (dose of 60 mg kg−1, Ketalar/Ketaminol vet 50 mg/ml; Pfizer Oy Animal Health, Espoo, Finland). The pupil of rats were dilated 10 minutes before IVT injections and eye imaging by applying 10 µl of topical tropicamide to each eye (Oftan tropicamid 5 mg/ml, Santen Pharmaceutical Co., Ltd., Tampere, Finland). Local ocular surface anesthesia was applied shortly before IVT injection by topical instillation of oxybuprocaine (Oftan® Obucain, 4 mg/mL; Santen Pharmaceutical Co., Ltd., Tampere, Finland). The IVT injection (injected volume of 1 μL) of 5 µM ML240-loaded mPEG_5kDa_-cholane and drug-free-mPEG_5kDa_-cholane nanosystems was performed using a 34 G needle connected to a syringe (Hamilton Co, Reno, NV, USA). Immediately after IVT injections and during imaging, the eyes were topically covered with topical carbomer hydrogel (Viscotears®, 2 mg/g; Dr. Winzer Pharma, Berlin, Germany) to prevent corneal dryness. After IVT injection and eye imaging, the animals were awakened by subcutaneous injection of atipamezole (Antisedan vet 5 mg/mL, Orion, Finland).

### Statistics

All data, unless otherwise indicated, were analyzed and graphed using Excel (Microsoft) and GraphPad Prism 7.05 for Windows. The threshold for statistical significance was identified as a P value of 0.05 or less. The TUNEL positive cells in the entire photoreceptor layer (outer nuclear layer-ONL) of four cross-sections per culture were counted manually (4-10 images from at least three retinae per treatment were taken and illustrated as open circles). The percentage of positive cells was calculated by dividing the absolute number of positive cells by the total number of cells in ONL (ONL area/the size of a photoreceptor nucleus (17.3 µm^2^)). The number of remaining photoreceptor cell rows in ONL was quantified by counting DAPI-stained cells in a linear row in 3 linear rows in four cross-sections per culture and averaging the counts. A one-way ANOVA test followed by Tukey’s multiple comparisons test was used to compare the different groups.

## 4. Results

VCP inhibitors include positively charged and polar chemical groups and hydrophobic residues [32] (Fig. 1A). The low aqueous solubility of VCP inhibitors requires their solubilization with DMSO, but this vehicle has safety concerns for therapeutic applications [33]. In addition, solutions of small molecules usually imply rapid elimination from the eye upon intravitreal administration [34]. Nanoformulations can simultaneously provide for both increased VCP inhibitor solubility and sustained release in the vitreous. To this aim, the VCP inhibitor ML240 was encapsulated in colloidal particles obtained with self-assembling mPEG_5kDa_-cholane and mPEG_5kDa_- cholesterol (Fig. 1 A-C).

**Figure 1:**
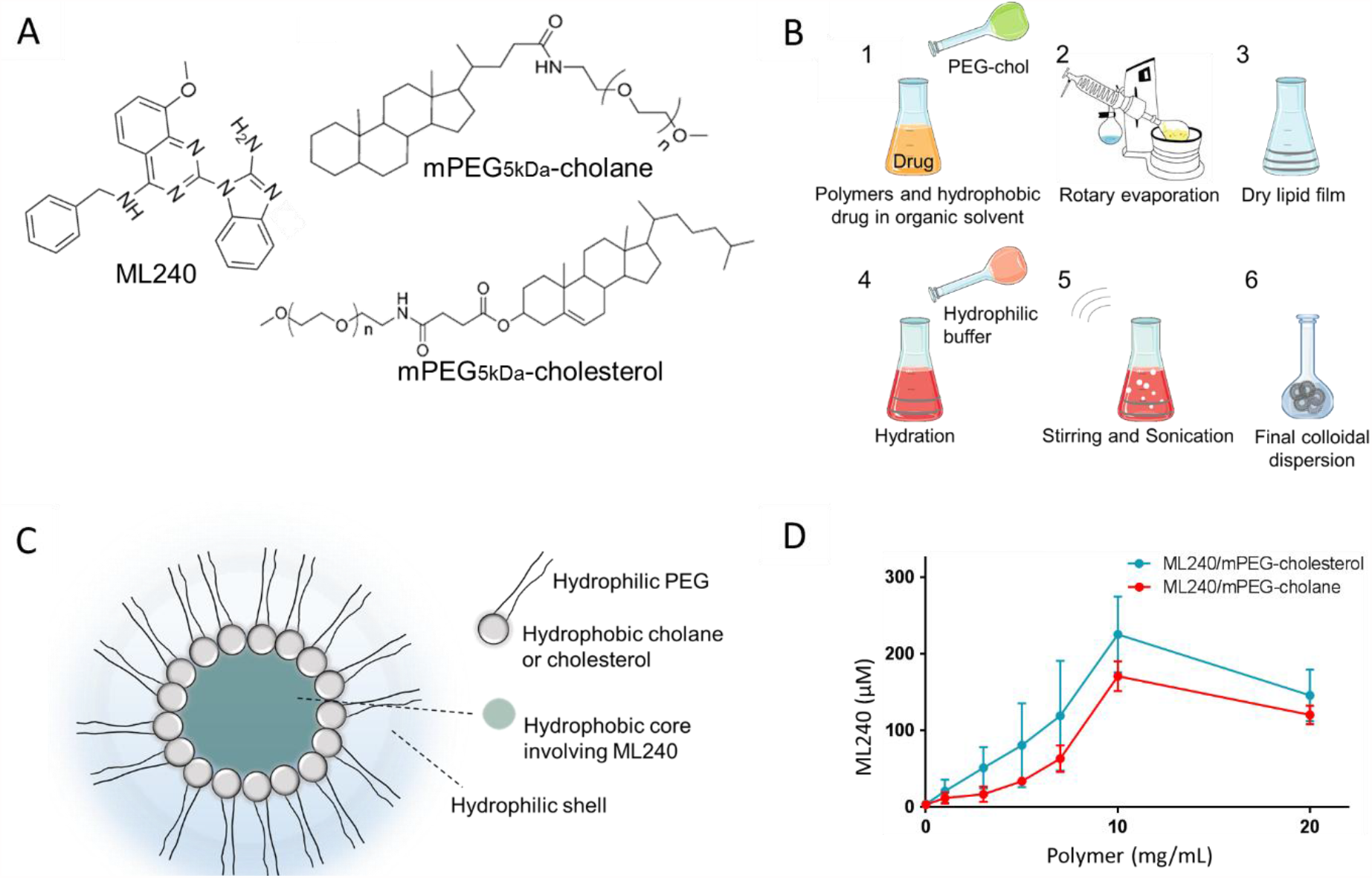
(A) Chemical structures of ML240, mPEG_5kDa_-cholane, and mPEG_5kDa_-cholesterol. (B) Schematic representation of the “film hydration” procedure to generate a colloidal dispersion by physical ML240 encapsulation in mPEG-coated particles (1-6). (C) Graphical representation of the self- assembled ML240 encapsulating system. (D) Solubility profile of ML240 at increasing concentrations of mPEG_5kDa_-cholane or mPEG_5kDa_-cholesterol polymers. The data are presented as mean ±SD (n=3).

### Encapsulation of the VCP inhibitor ML240 in colloidal mPEG_5kDa_-cholane and mPEG_5kDa_-cholesterol systems

The efficiency of physical encapsulation and dispersibility of ML240 in aqueous medium was investigated by assessing the drug dissolved in the aqueous medium with increasing mPEG_5kDa_-cholane or mPEG_5kDa_-cholesterol concentrations and constant excess of fed drug (0-10 polymer/drug weight ratios). The results (Fig. 1D) show that without mPEG_5kDa_-cholane or mPEG_5kDa_-cholesterol, the ML240 solubility was negligible (3 μM), while both mPEG_5kDa_-cholane and mPEG_5kDa_-cholesterol increased the ML240 colloidal dispersion to yield maximal drug solubility with 10 mg/mL of mPEG_5kDa_-cholane and mPEG_5kDa_-cholesterol: 170.6±19.5 µM and 225.3±9.4 µM, respectively. A slight concentration decrease was observed at higher polymer concentrations (20 mg/mL), as already observed with other drugs [24].

Figure 2A shows that the size of the ML240 loaded mPEG_5kDa_-cholane and mPEG_5kDa_- cholesterol nanoparticles were 54.8 ± 8.4 nm and 31.9 ± 3.1 nm, respectively. The polydispersity indices were 0.38 ± 0.07 and 0.37 ± 0.05 for the mPEG_5kDa_-cholane and mPEG_5kDa_-cholesterol nanoparticles, respectively. The results were confirmed by TEM analyses showing spherical particles with some size heterogeneity (Fig. 2C-D). The size of the nanoparticles remained fairly stable for 72 hours in buffer and porcine vitreous (Fig. 2A-B).

**Figure 2:**
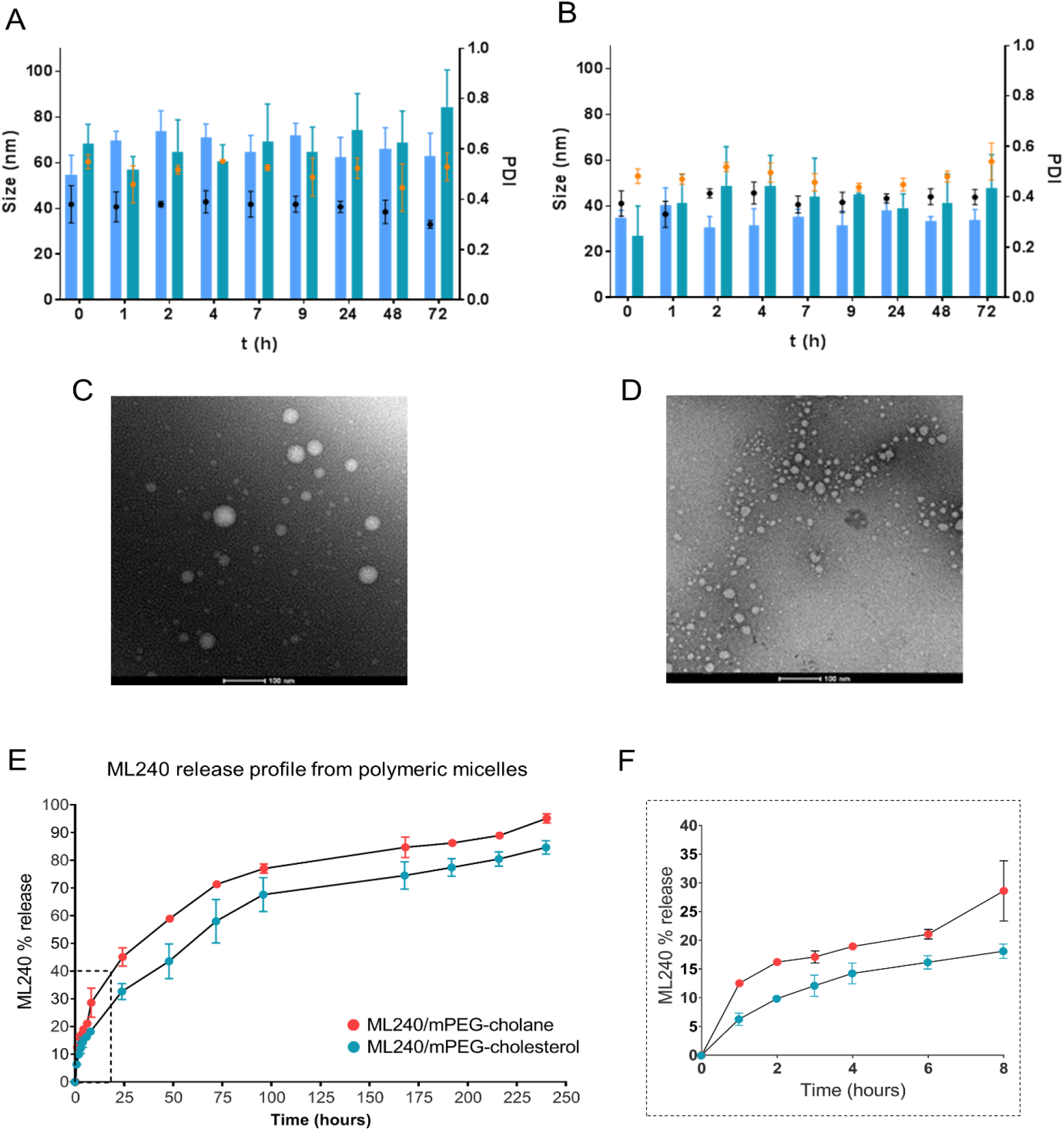
Characterization of ML420-loaded mPEG_5kDa_-cholane and mPEG_5kDa_-cholesterol nanoparticles. Mean size and PDI of ML240/mPEG_5kDa_-cholane (A) and ML240/mPEG_5kDa_-cholesterol (B) particles in PBS, pH 7.4 (size: ▪ PDI •) and in porcine vitreous (size ▪; PDI •), respectively. TEM images of ML240/mPEG_5kDa_-cholane (C) and ML240/mPEG_5kDa_-cholesterol (D) nanoparticles. (E-F) ML240 release from mPEG_5kDa_-cholane and mPEG_5kDa_-cholesterol nanoparticles in PBS, pH 7.4. (F) Magnification of the initial release profile from panel E. The data are presented as mean ±SD (n=3). PDI: polydispersity index.

*In vitro* drug release profiles are shown in Figure 2E-F. ML240 was released from the mPEG_5kDa_-cholane and mPEG_5kDa_-cholesterol nanoparticles over 10 days. Both formulations showed initial burst release during the first 8 hours, 17% and 28% drug release for mPEG_5kDa_-cholane and mPEG_5kDa_-cholesterol nanoparticles, respectively. Thereafter, the drug was released slowly, achieving 88-98% release in 10 days.

### Effects of ML240-loaded nanoparticles on photoreceptor cell survival in organotypic RHO^P23H^ rat retinal cultures

The neuroprotective effects and possible functional alterations in retinal organotypic cultures were investigated after exposure to ML240 nanoformulated with mPEG_5kDa_- cholane and mPEG_5kDa_-cholesterol. We used RHO^P23H^ retinae isolated at PN9 and cultured for 6 days on polycarbonate membrane inserts. At this time, PN15, photoreceptor cell death in culture mimics the *in vivo* peak of degeneration of the RHO^P23H^ rat retinal model (Fig. 3A, [11,35]). We applied 5 µM ML240 concentration of each formulation or the equivalent concentration of drug-free mPEG_5kDa_-cholane and mPEG_5kDa_-cholesterol to the culture medium and replaced the culture medium and treated the explants every two days (Fig. 3B). We also compared a therapeutic concentration provided by the drug alone (20 µM ML240 dissolved in DMSO) as reported previously in the same animal model [11]. Untreated RHO^P23H^ retinal explants were used to assess the baseline phenotype. Neuroprotective effects were assessed using the TUNEL assay (Fig. 3C). The percentage of TUNEL-positive cells in the outer nuclear layer (ONL) of untreated RHO^P23H^ retinae was 4.8 ± 1.6 %, and cell death was further increased to 6.1 ± 2.1 % in DMSO-treated retinae (Fig. 3D). Nanoformulations of ML240 allowed to overcome the intrinsic toxicity of DMSO, as the ONL cell death in RHO^P23H^ retinae decreased to 3.9 ± 0.7 % (ML240 formulated with mPEG_5kDa_-cholane) and 3.7 ± 0.5 % (ML240 formulated with mPEG_5kDa_-cholesterol). These levels were significantly (**p<0.01) smaller than the DMSO-treated retinae. ML240-free polymeric nanosystems showed similar TUNEL staining as the untreated controls, confirming the safety of the polymers.

**Figure 3:**
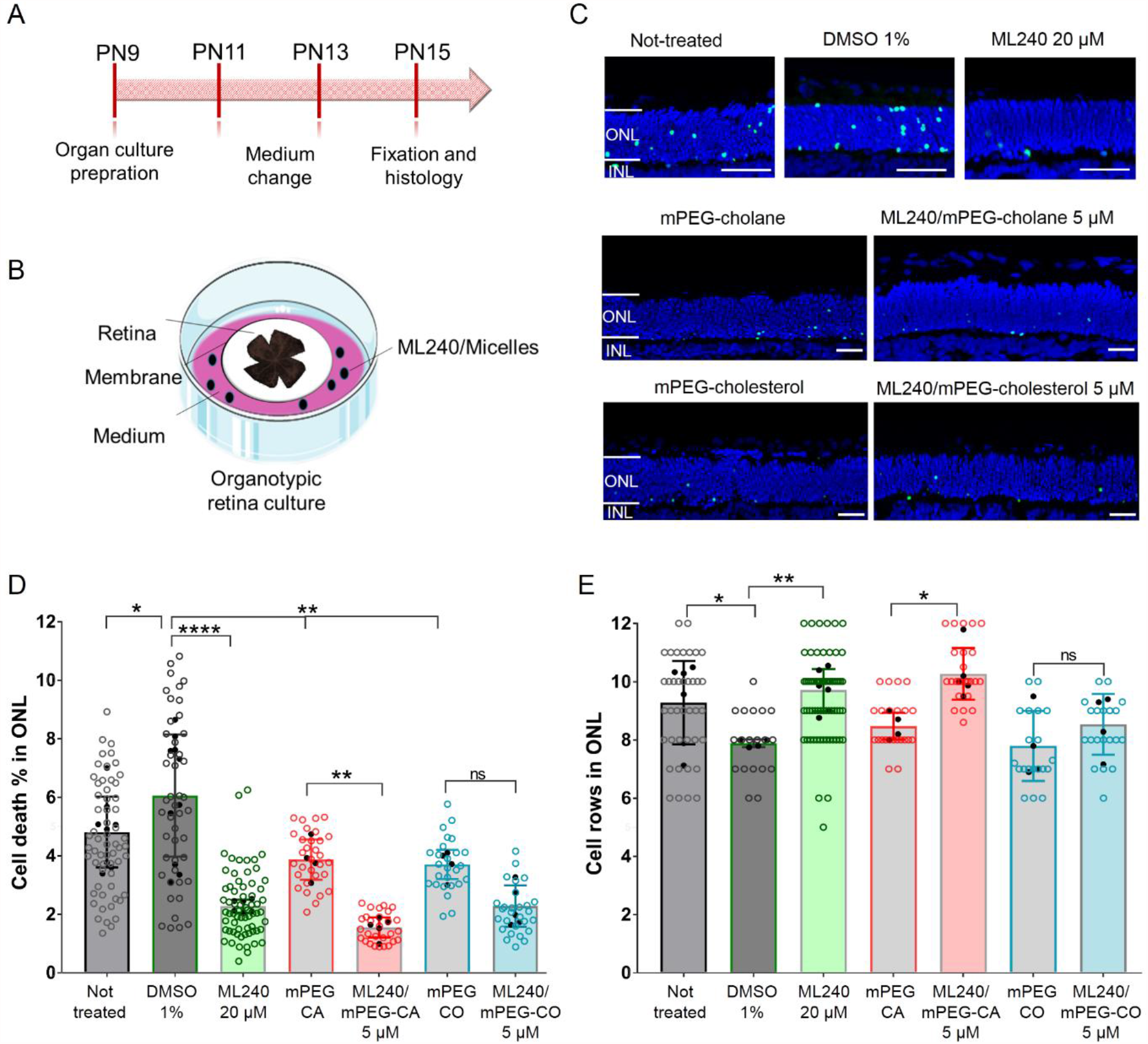
Delivery of ML240 by mPEG_5kDa_-cholane nanoparticles enhances photoreceptor cell survival in organotypic RHO^P23H^ rat retinal cultures. (A) Timeline of photoreceptor degeneration in RHO^P23H^ heterozygous transgenic rats and experimental setup. Retinae from RHO^P23H^ transgenic rats were explanted at PN day 9, cultivated for 6 days (PN9 DIV6), and treated every second day. (B) Schematic representation of the interphase organotypic retinal culture. (C) Explants were stained with the TUNEL assay to differentiate photoreceptors undergoing cell death (green) using nuclei counterstaining with DAPI (blue). (D) Bar chart shows the percentage of TUNEL-positive cells in the ONL (E) Quantification of the number of photoreceptor cell rows in the ONL. Scale bar is 50 µm. Open circles represent the evaluated images. Closed circles represent the n value. Data plotted as mean ± SD. One-way ANOVA analysis was performed at ****p<0.0001, **p<0.01, *p<0.05. Abbreviations: ONL: outer nuclear layer, INL: inner nuclear layer, CA: cholane, CO: cholesterol.

We found lower cell death after exposure to 20 µM ML240, 5 µM ML240-loaded mPEG_5kDa_-cholane and mPEG_5kDa_-cholesterol nanoparticles than in the untreated control, thereby showing the neuroprotective effect of VCP inhibition in RHO^P23H^ rat retinae (Fig. 3C). A four times lower dose of ML240 formulated with mPEG_5kDa_-cholane was sufficient to achieve a comparable photoreceptor protection to the drug alone (cell death for 20 µM ML240 alone: 2.3 % ± 0.2; cell death for 5 µM ML240 in mPEG_5kDa_- cholane nanoparticles: 1.6 % ± 0.3) (Fig. 3D). A significant decrease of cell death in retinae treated with 5 µM ML240-loaded mPEG_5kDa_-cholane nanoparticles compared to ML240-free mPEG_5kDa_-cholane nanosystems was observed (cell death for 5 µM ML240-loaded mPEG_5kDa_-cholane nanoparticles: 1.6 % ± 0.3; cell death for ML240- free mPEG_5kDa_-cholane nanosystems: 3.9 % ± 0.7, **p<0.01). However, the cell protection effect was not reflected in RHO^P23H^ retinae treated with ML240-loaded mPEG_5kDa_-cholesterol nanoparticles and ML240-free mPEG_5kDa_-cholesterol nanosystems (cell death for 5 µM ML240-loaded mPEG_5kDa_-cholesterol nanoparticles: 2.3 % ± 0.7; cell death for ML240-free mPEG_5kDa_-cholesterol nanosystems: 3.7 % ± 0.5).

Photoreceptor cell survival was derived from the count of cell nuclei of DAPI stained photoreceptors in different sections of the treated retinae. As expected, the RHO^P23H^ retinae treated with 20 µM ML240 displayed significantly more photoreceptor cell rows compared to the DMSO control (cell rows for 20 µM ML240: 9.7 ± 0.7; cell rows for DMSO/cell culture medium: 7.9 ± 0.1, **p<0.01) (Fig. 3E). However, the incubation with only DMSO/cell culture medium reduced photoreceptor cell survival compared to the untreated control. Furthermore, we also found that the RHO^P23H^ retinae treated with ML240 formulated with polymeric nanoparticles showed significantly higher photoreceptor cell survival when compared to the untreated control (cell rows for 5 µM ML240-loaded mPEG_5kDa_-cholane nanoparticles: 10.3 ± 0.9; cell rows for ML240-free mPEG_5kDa_-cholane nanosystems: 8.5 ± 0.5, *p<0.05; cell rows for 5 µM ML240-loaded mPEG_5kDa_-cholesterol nanoparticles: 8.5 ± 1.0: cell rows for ML240-free mPEG_5kDa_- cholesterol nanosystems: 7.8 ± 1.2; cell rows for untreated: 9.3 ± 1.4). However, only ML240-loaded mPEG_5kDa_-cholane nanoparticles showed significant improvement of photoreceptor cell rows compared to the drug-free control particles, and we could not see this effect when comparing ML240-loaded mPEG_5kDa_-cholesterol nanoparticles and drug-free nanosystems control. There was no significant difference between RHO^P23H^ retinal explants treated with the 20 µM ML240 alone and 5 µM ML240-loaded mPEG_5kDa_-cholane nanoparticles, which reflected a similar behavior that was observed for the recovery of degenerated photoreceptors in the ONL. Retinae exposed to ML240-free polymeric nanosystems showed no difference in photoreceptor cell survival compared to the untreated control, confirming that the formulations themselves did not affect retinal morphology.

### RHO^P23H^ rat retinae treated with ML240-loaded nanoparticles restored rhodopsin distribution in vitro

The P23H mutation causes rhodopsin mislocalization, leading to photoreceptor degeneration, and defects in rhodopsin transport are associated with disorganized and shortened outer segments [36]. Restoration of rhodopsin in the outer segment is important for proper rhodopsin function and is crucial for reversing the adRP phenotype. To test the functional effect of nanoformulated ML240, we examined rhodopsin localization in RHO^P23H^ rat retinal explants. While wild-type rhodopsin is mainly localized in the outer segments of rod photoreceptors, P23H mutated rhodopsin is abnormally distributed throughout the inner part of the ONL (Fig. 4A). Also, RHO^P23H^ rat retinal explants treated with drug-free polymeric nanosystems displayed larger amounts of rhodopsin in the ONL (Fig. 4B, left panels).

**Figure 4:**
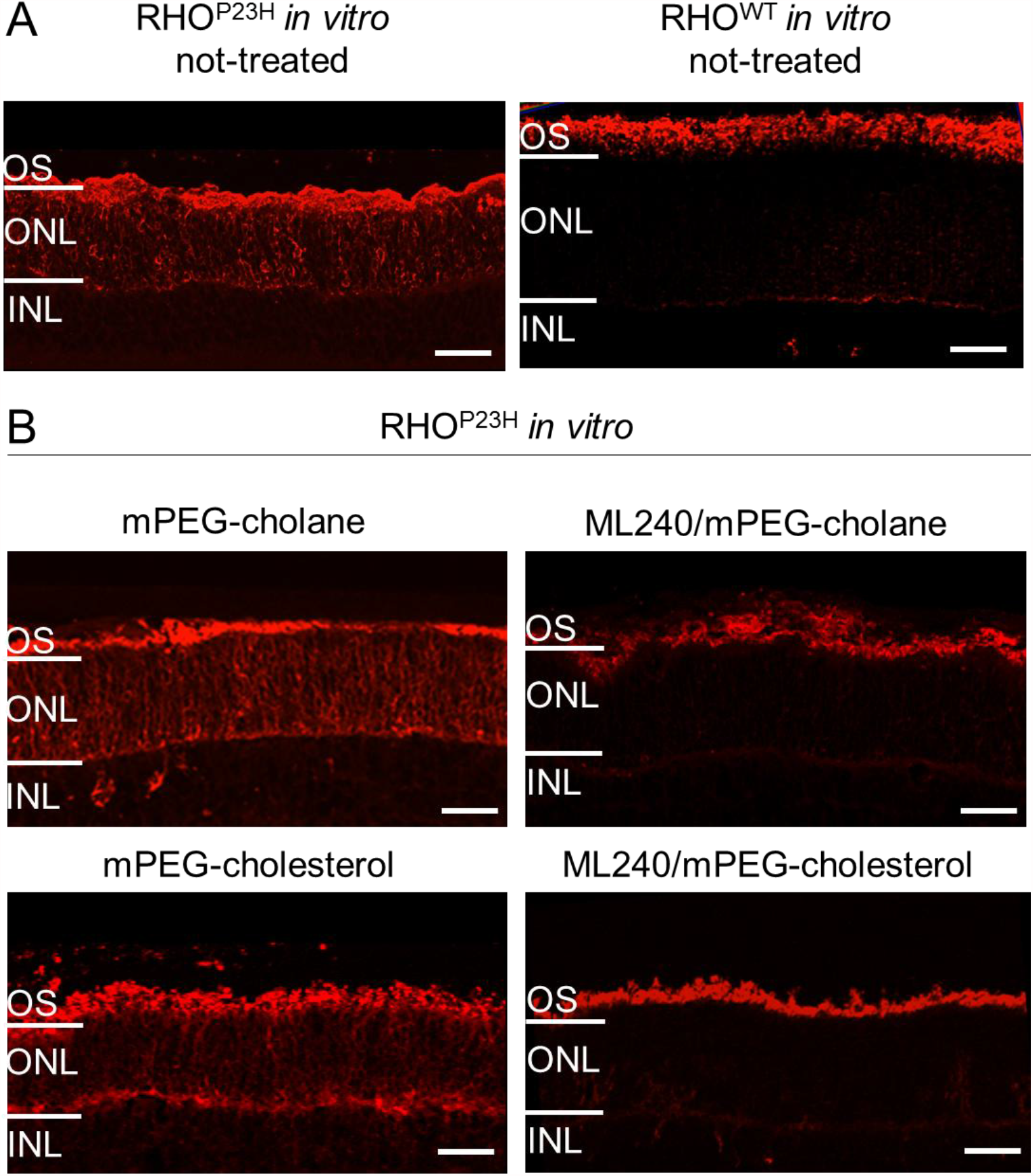
RHO^P23H^ retinae treated with ML240-loaded polymeric nanoparticles restored rhodopsin distribution (red immunostaining) *in vitro*. (A) Not treated retinae from RHO^P23H^ and RHO^WT^ rats explanted at PN9 and cultivated for 6 days. (B) Retinae from RHO^P23H^ treated every second day with 5 µM ML240-loaded mPEG_5kDa_-cholane nanoparticles or 5 µM ML240-loaded mPEG_5kDa_-cholesterol nanoparticles. Corresponding controls were treated using the same volume of the appropriate vehicle. Scale bar is 50 µm.

Immunohistochemistry confirmed that RHO^P23H^ retinal cultures treated with ML240 encapsulated in mPEG_5kDa_-cholane or mPEG_5kDa_-cholesterol nanoparticles showed enrichment in the rhodopsin localization in the outer segment as in the wild-type phenotype (Fig. 4A and 4B, right panels).

In conclusion, treatment with ML240-loaded mPEG_5kDa_-cholane or mPEG_5kDa_- cholesterol nanoparticles restores proper rhodopsin localization to the outer segment.

### Effects of ML240-loaded nanoparticles on inflammation in organotypic RHO^P23H^ rat retinal cultures

Next, we tested the anti-inflammatory effects of VCP inhibition in the *RHO*^P23H^ retinae by staining with an antibody against ionized calcium-binding adapter molecule 1 (Iba- 1). Iba1 levels increase during microglial activation, an important event in the inflammatory response pathway of the retina.

In their resting state, as in RHO^WT^, microglial cells are localized only in the inner plexiform layer (IPL) of the retina and in the outer plexiform layer (OPL) from where they extend their projections across the different nuclear layers (Fig. 5A). In their active state, as observed in the mutated retinae of RHO^P23H^ rats, microglial cells are recruited and migrate through the photoreceptor layer (ONL) to promote inflammation [37,38] (Fig. 5A).

**Figure 5:**
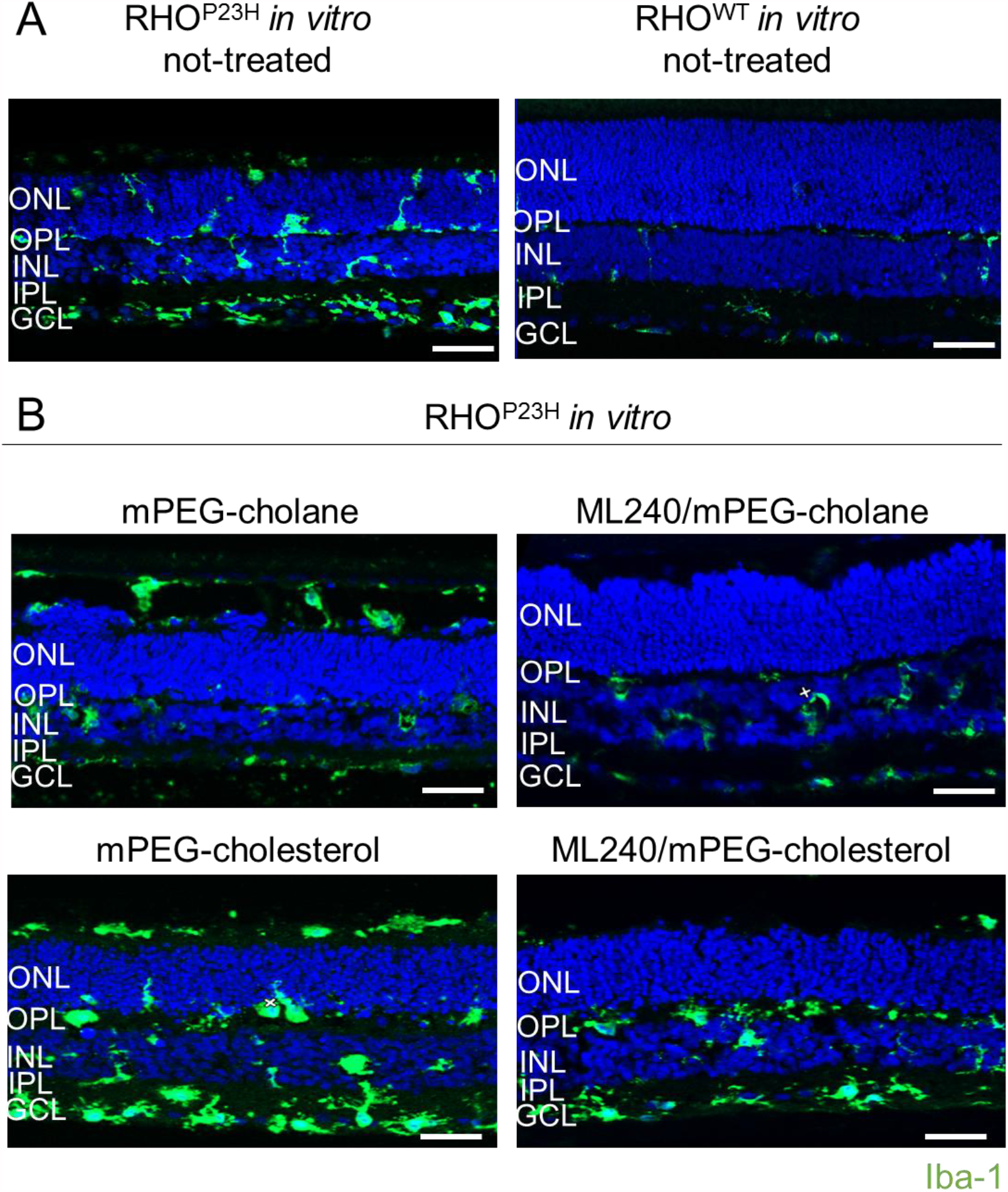
Immunostaining of Iba1 (green) indicates reduced microglial activation in RHO^P23H^ retinal cultures treated with 5 µM ML240-loaded mPEG_5kDa_-cholane nanoparticles than cultures treated with 5 µM ML240-loaded mPEG_5kDa_-cholesterol nanoparticles. (A) Not treated retinae from RHO^P23H^ and RHO^WT^ rats explanted at PN9 and cultivated for 6 days. (B) Retinae from RHO^P23H^ treated every second day with 5 µM ML240-loaded mPEG_5kDa_-cholane nanoparticles or 5 µM ML240-loaded mPEG_5kDa_- cholesterol nanoparticles. Corresponding controls were treated using the same volume of the appropriate vehicle. Scale bar is 50 µm.

Immunostaining of Iba1 treated with ML240-free mPEG_5kDa_-cholane or mPEG_5kDa_-cholesterol nanosystems showed positive Iba1 staining mainly in the IPL and OPL with some Iba1-positive areas in the GCL, in the ONL and INL (Fig. 5B). Retina treated with ML240-loaded mPEG_5kDa_-cholesterol nanoparticles presented higher microglial inflammation-related activation than those treated with ML240-loaded mPEG_5kDa_- cholane nanoparticles, as indicated by an increase in the number of microglial cells migrating from the GCL to the adjacent layers (Fig 5B). We observed less migration of Iba1-positive cells to the ONL of retina treated with ML240-loaded mPEG_5kDa_- cholane nanoparticles than ML240-loaded mPEG_5kDa_-cholesterol nanoparticles treated retinae (Fig. 5B). The lesser apical projections of microglial cells in ML240-loaded nanoparticles treated retinae relative to their drug-free controls showed that VCP inhibition can suppress microglial activation that goes along with the P23H mutation.

Taken together, our data indicate that efficient encapsulation of VCP inhibitor delivered by mPEG_5kDa_-cholane nanoparticles suppresses microglial inflammation in RHO^P23H^ retinal explants. We hence focused on ML240-loaded mPEG_5kDa_-cholane nanoparticles for their short-term and long-term efficacy studies in RHO^P23H^ rat retinal explants by mimicking IVT.

### ML240 formulated with mPEG_5kDa_-cholane enables long-term photoreceptor cell survival in RHO^P23H^ retinal explants

Next, we prepared organotypic retinal cultures from RHO^P23H^ rats at PN9 and incubated these cultures for 6 days (up to PN15), as short-term treatment model, and 21 days (up to PN30), as long-term treatment model, by administering the formulations from the GCL side of the retina (Fig. S1). Again, TUNEL assays were used.

The RHO^P23H^ retinae treated with ML240-loaded mPEG_5kDa_-cholane nanoparticles showed a significant reduction of the percentage of cell death in the ONL after 6 days treatment (short-term) compared with the untreated and ML240-free mPEG_5kDa_- cholane nanosystems treated control (cell death for ML240-loaded mPEG_5kDa_-cholane nanoparticles: 0.93 % ± 0.05, ***p<0.001 to untreated retinae, *p<0.05 to drug-free mPEG_5kDa_-cholane nanosystems; cell death for mPEG_5kDa_-cholane nanoparticles: 1.95 % ± 0.29; cell death for untreated: 3.10 % ± 0.41) (Fig. 6A, 6C). Interestingly, the drug-free mPEG_5kDa_-cholane nanosystems also showed a lower cell death rate than the untreated group, which needs further investigation. In addition, cultures treated with ML240-loaded mPEG_5kDa_-cholane nanoparticles led to significant increase in surviving photoreceptor cell rows in the ONL (cell rows for ML240-loaded mPEG_5kDa_-cholane nanoparticles treatment: 10.38 ± 0.12, ***p<0.001 to untreated retinae, *p<0.05 to drug-free mPEG_5kDa_-cholane nanosystems; cell rows for drug-free mPEG_5kDa_-cholane nanosystems treatment: 8.11 ± 0.46; cell rows for untreated: 7.34 ± 0.74) (Fig. 6A, 6D).

**Figure 6:**
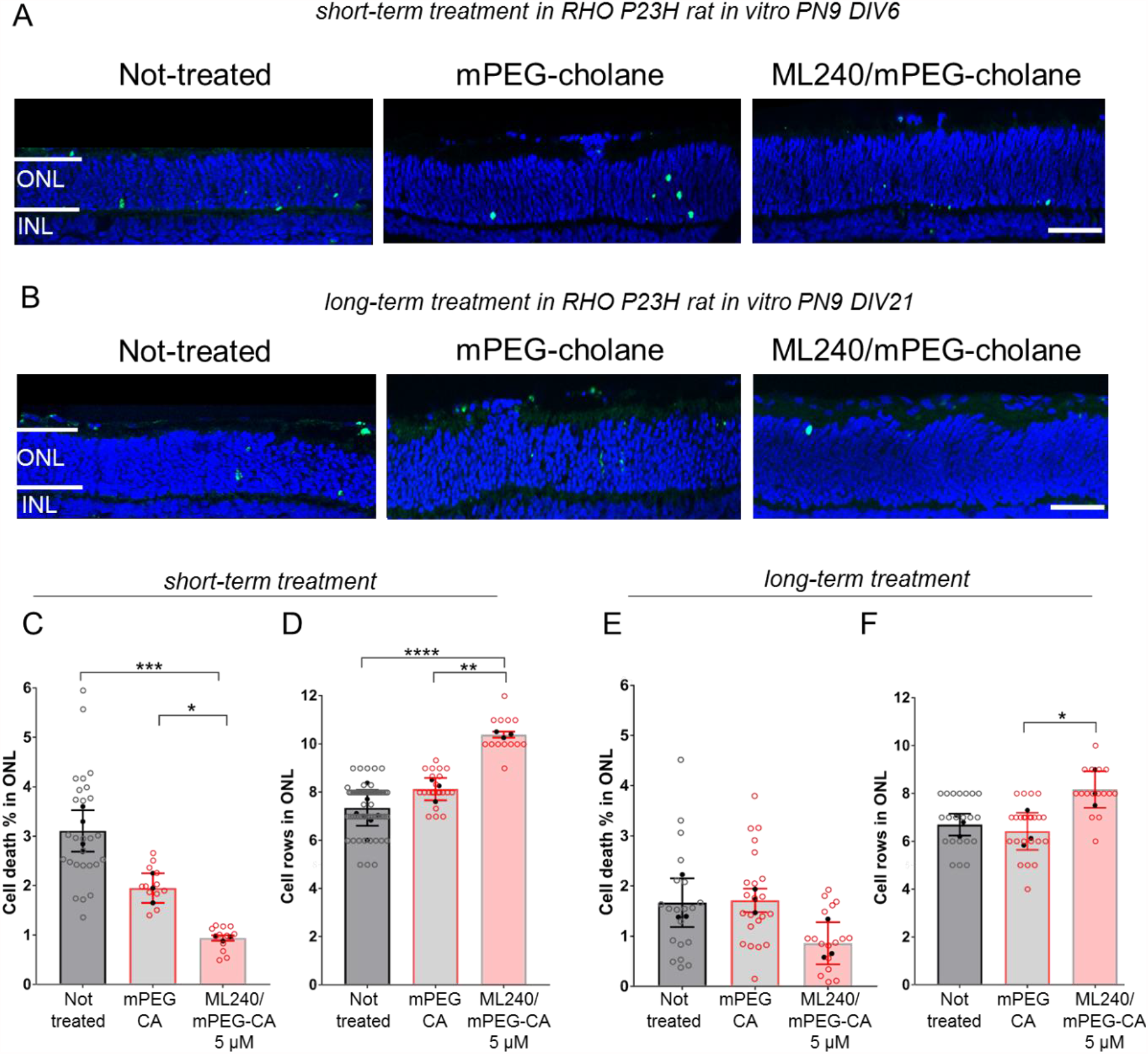
Enhanced photoreceptor cell survival in RHO^P23H^ retinae treated with ML240-loaded mPEG_5kDa_-cholane nanoparticles after (A) short- (6 days) and (B) long-term (21 days) treatments by administering the formulations from the GCL side of the retina. TUNEL staining of undergoing cell death (green), nuclei counterstaining with DAPI (blue). Scale bar is 50 µm. (C) Bar chart shows the percentage of TUNEL-positive cells in the ONL after short-term treatment. (D) Quantification of the number of cell rows in the ONL after short-term treatment. (E) Bar chart shows the percentage of TUNEL-positive cells in the ONL after long-term treatment. (F) Quantification of the number of cell rows in ONL after long-term treatment. Open circles represent the evaluated images. Closed circles represent the n value. Data plotted as mean ± SD. One-way ANOVA analysis was performed at ****p<0.0001, ***p<0.001, **p<0.01, *p<0.05. Abbreviations: PN: postnatal day, DIV: days *in* vitro, mPEG-CA: monomethoxy-PEG_5kDa_-cholane.

The neuroprotective effect of 5 µM ML240-loaded mPEG_5kDa_-cholane nanoparticles was also observed after 21 days of treatment (long-term). RHO^P23H^ rat retinae were explanted at PN9 and cultured up to PN30 (Fig. S1, Fig. 6B, Fig. 6E-F). The ML240- loaded mPEG_5kDa_-cholane nanoparticles treatment resulted in a significantly higher number of surviving photoreceptor cells in the ONL (long-term treatment; surviving photoreceptor cell rows for ML240-loaded mPEG_5kDa_-cholane nanoparticle treatment: 8.16 ± 0.76, *p<0.05 to drug-free mPEG_5kDa_-cholane nanosystems; surviving photoreceptor cell rows for drug-free mPEG_5kDa_-cholane nanosystems: 6.41 ± 0.77; surviving photoreceptor cell rows for untreated: 6.69 ± 0.45) (Fig. 6F). We observed a decrease in cell death in photoreceptors after 21 days of treatment, but it was not significant between untreated, drug-free mPEG_5kDa_-cholane nanosystems and drug- loaded nanoparticles treated retinal cultures (Fig. 6E), which was expected for the long-term cultures, as the retina has no blood supply and undergoes axotomy-induced stress responses during long-term culturing [39].

To examine possible side effects of ML240-loaded mPEG_5kDa_-cholane nanoparticles, we intravitreally injected them into wild-type adult rat eyes (Fig. 7). This was done in comparison to a drug-free mPEG_5kDa_-cholane control nanosystem. Eventual adverse effects as a consequence of iatrogenic injury during IVT were quality controlled by fundus camera and OCT in rats before and two weeks after injection (Fig. 7, Fig S2). Long-term IVT administration showed good tolerability of the formulations without any visible alterations of the fundus such as conjunctival bleeding, induction or worsening of cataracts, retinal detachment, or other retinal morphological alterations (Fig. 7A, Fig S2). There were no signs of vitreous floaters or infiltrated cells in the vitreous after two weeks. There was a slight increase in the retinal vascular tortuosity in both formulations two weeks after injections. No activation of the cell death was detected by examining TUNEL assay and counting photoreceptor cell rows (Fig. 7 B-C). This supports the safety of the ML240-loaded mPEG_5kDa_-cholane formulation as well as the good tolerability of the drug-free polymeric nanosystem itself.

**Figure 7:**
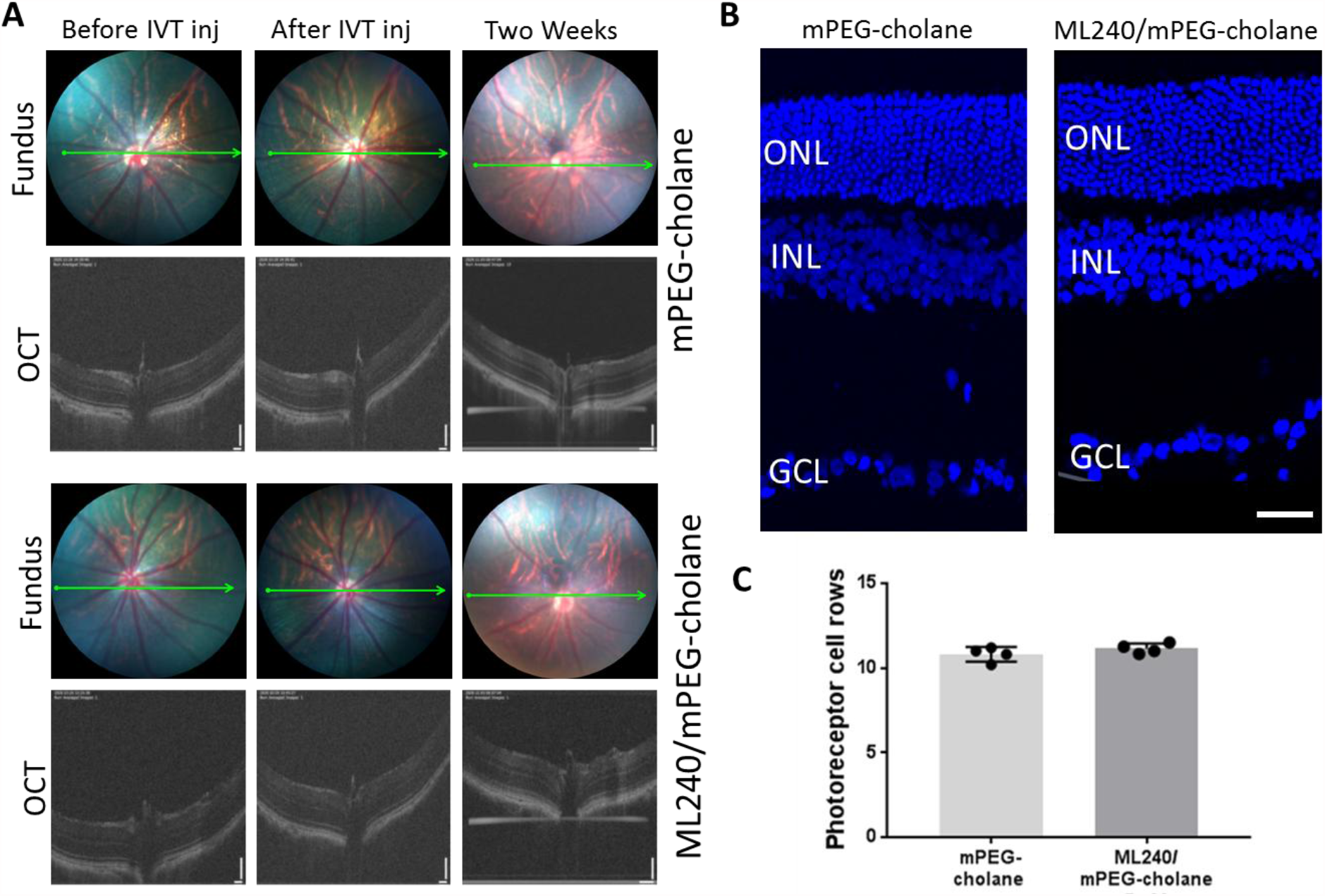
Optical Coherence Tomography (OCT) and fundus images of rat vitreous/retina before IVT injection (inj), immediately after IVT injection and following two weeks after IVT injection of 5 µM ML240-loaded mPEG_5kDa_-cholane nanoparticles and free-drug mPEG_5kDa_-cholane nanosystems (A). Quantification of the number of photoreceptor cell rows in ONL by counting DAPI-stained cells in a linear row in 3 linear rows in 3 different vertical sections (B-C).

## 5. Discussion

Previous findings suggest that VCP inhibition may have therapeutic value in the treatment of photoreceptor cell loss and adRP [10,11]. However, VCP inhibitors usually have poor solubility, limiting their pharmaceutical exploitation and delivery to the eye. Here, we explored an approach to efficiently enhance the solubility of a hydrophobic VCP inhibitor by physical encapsulation in self-assembling mPEG-polycyclic based nanoparticles. We show that this approach improves the solubility of the VCP inhibitor ML240 by two orders of magnitude. In addition, we demonstrate the sustained release of ML240 from the nanoparticles. Notably, the VCP inhibitor delivered by these polymer-based nanoparticles showed superior short-and long-term neuroprotection in RHO^P23H^ rat retinal explants.

Self-assembling amphiphilic polymers can be successfully exploited to enhance the solubility of drugs by simple, reproducible, and scalable processes. Moreover, due to their small size, polymer-based nanoparticles are capable of diffusing through relevant tissues and biological matrices. Previously, we showed that nanoparticles of 50-200 nm in diameter were diffusing in the vitreous [40] and PEG-coated liposomes of the size of ≈50 nm penetrated across the inner limiting membrane in the *ex vivo* bovine eyes [19]. Thus, it is likely that the nanoparticles in this study are capable of permeating in the vitreous and retina.

In the study reported here, mPEG_5kDa_-cholane and mPEG_5kDa_-cholesterol were used to encapsulate the VCP inhibitor ML240. These amphiphilic polymers have similar polycyclic functional groups at one end of the mPEG chain (Fig. 1A), although the polycyclic groups in these two compounds are inverted [24]. The different orientations of the polycyclic moieties provide different structural arrangements of the hydrophobic moiety when mPEG_5kDa_-cholane and mPEG_5kDa_-cholesterol self-assemble into colloidal systems [26], resulting in different drug encapsulation and stability profiles of the resulting nanosystems [24].

Both nanoparticles obtained with the amphiphilic polymers remarkably improved the aqueous dispersion of ML240. The nanoparticles obtained with mPEG_5kDa_-cholane were about 70% bigger than those obtained with mPEG_5kDa_-cholesterol, suggesting that the nanoparticle structure could be significantly different for the two formulations. Importantly, the formulations were colloidally stable in vitreous up to 72 hours, indicating that the amphiphilic polymer coating of the drug core can prevent aggregation that could be driven by the adsorption of the vitreous components, namely collagen and hyaluronic acid, as observed with other systems [34].

Drug release from mPEG_5kDa_-cholane and mPEG_5kDa_-cholesterol nanoparticles showed similarities. Both systems showed a burst release for about 8 hours, followed by the slow release in about 10 days. However, the burst and release rates with the mPEG_5kDa_-cholane nanoparticles were higher than those with the mPEG_5kDa_-cholesterol nanoparticles. These results may be due to different orientations of the drug and polymer in the nanoparticles, possibly due to different interactions between the polycyclic moiety of the polymers with the drug, resulting in more unstable structures in the case of mPEG_5kDa_-cholane nanoparticles compared to the mPEG_5kDa_-cholesterol formulation.

The delivery of ML240 formulated with mPEG_5kDa_-cholane resulted in improved drug efficacy over ML240 dissolved in DMSO: lower cell death rate and higher photoreceptor cell survival. This may be due to the sustained release of the drug exerting its therapeutic activity to the target cells gradually while avoiding the peak concentration and fast diffusion kinetics of the soluble free drug. On the other hand, the nanoparticles may also facilitate drug transfer into the retina. An efficient dispersion and delivery were achieved with the nanoformulated drug, resulting in an equally potent effect (increased photoreceptors survival) using only 25% of the dose required with the unformulated drug. The ML240 concentration used *in vitro* to achieve a therapeutic effect by encapsulating the drug in polymeric nanoparticles, namely 5 µM, showed the same neuroprotective effect as the unformulated drug (20 µM in DMSO) [11]. The lower toxicity of mPEG_5kDa_-cholane also facilitates the overall therapeutic activity as compared with DMSO.

The treatment with ML240-loaded mPEG_5kDa_-cholane nanoparticles resulted in a significant reduction in inflammatory microglial activation. However, this was not seen with ML240-loaded mPEG_5kDa_-cholesterol nanoparticles. The difference may be attributed to drug release kinetics or intracellular drug delivery. Furthermore, the higher toxicity of ML240-loaded mPEG_5kDa_-cholesterol nanoparticles compared to ML240 loaded mPEG_5kDa_-cholane nanoparticles may be due to different interactions of the mPEG-polycyclic component interactions with the cell membrane [24]. Cholesterol in the cell membranes may show asymmetric localization and diffusion [41]. Plasma membranes of the rod outer segments are rich in cholesterol and saturated fatty acids [42,43], making them sensitive to remodeling by membrane-associated lipids [42,44]. Thus, asymmetric localization and diffusivity of cholesterol in the cell membranes may be affected by the associated mPEG_5kDa_-cholesterol, resulting in altered membrane asymmetry, -heterogeneity, or -stability, ultimately triggering inflammation.

ML240-containing mPEG_5kDa_-cholane nanoparticles were selected to investigate for short- and long-term efficiency by mimicking IVT based on their superior performance. A 21 days’ RHO^P23H^ culture treatment with ML240-loaded mPEG_5kDa_-cholane nanoparticles resulted in significant photoreceptor protection and increased their survival. Again, we have reasons to believe that the sustained and prolonged release of the drug as a result of the encapsulation and local and sustained action of the VCP inhibitor likely produced this superior outcome. In the future, such formulations may thus allow intravitreal administration intervals similar to those used for anti-VEGF therapy to treat the wet form of AMD [45] and therefore be transferable to a routine clinical therapeutic regime.

As a perspective beyond drug-based VCP inhibition, our study may be considered proof of concept that a colloidal self-assembling system can serve as an advanced delivery system for treating retinal degeneration. Polymer-based nanoparticles can provide increased solubility, slow-release, and prolonged action of a drug combined with excellent ocular tolerability. Beyond the scope of our study, and as the retina is part of the central nervous system (CNS), we also suggest this technology for further studies to facilitate drug delivery to the CNS.

## Acknowledgments

This study was supported by funds (to M.Ue. and B.A-G) from FFB (Grant PPA-0717- 0719-RAD), the Kerstan Foundation, European Union’s Horizon 2020 research and innovation program under the Marie Skłodowska-Curie (Grant agreement No. 722717 – project OCUTHER), the Maloch Stiftung, and the ProRetina Foundation. The animal husbandry personnel at the Universitätsklinikums Tübingen and Norman Rieger are acknowledged for animal care.

## Supplementary files

**Figure S1:**
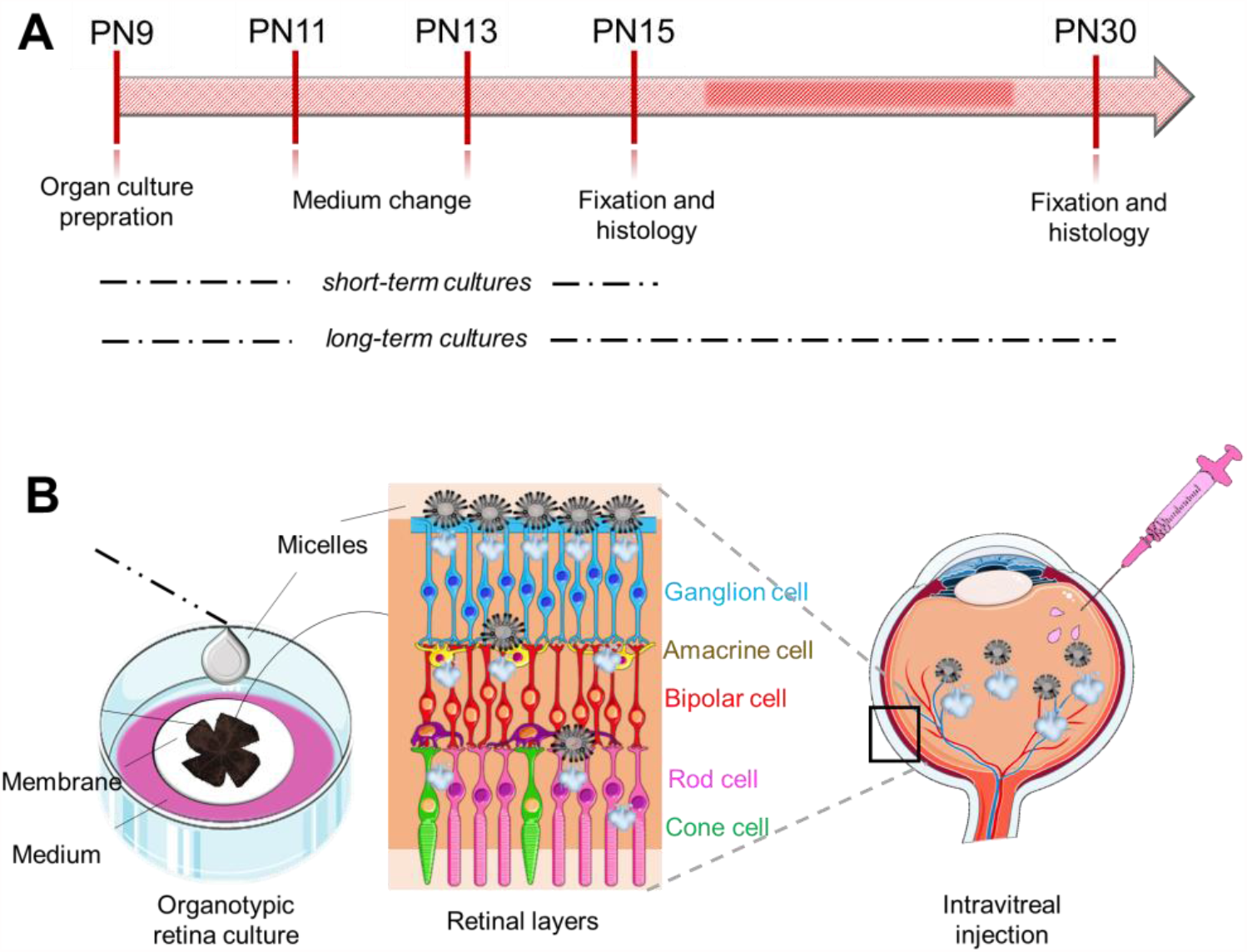
Experimental setup and schematic representation of the RHO^P23H^ retinal organotypic cultures by mimicking IVT. (A) The retinal cultures of RHO^P23H^ rodents at PN9 were prepared and incubated for 6 days (up to PN15) for short-term treatment and 21 days (up to PN30) for long-term treatment. The medium was changed every second day. (B) The retinae were dissected and placed on membranes with photoreceptors facing down. Treatment was performed directly on the retina by applying 5 µM ML240-loaded mPEG_5kDa_-cholane nanoparticles, and mPEG_5kDa_-cholane control vehicle once on the GCL layer. Untreated explants were cultured with medium alone and used as controls.

**Figure S2:**
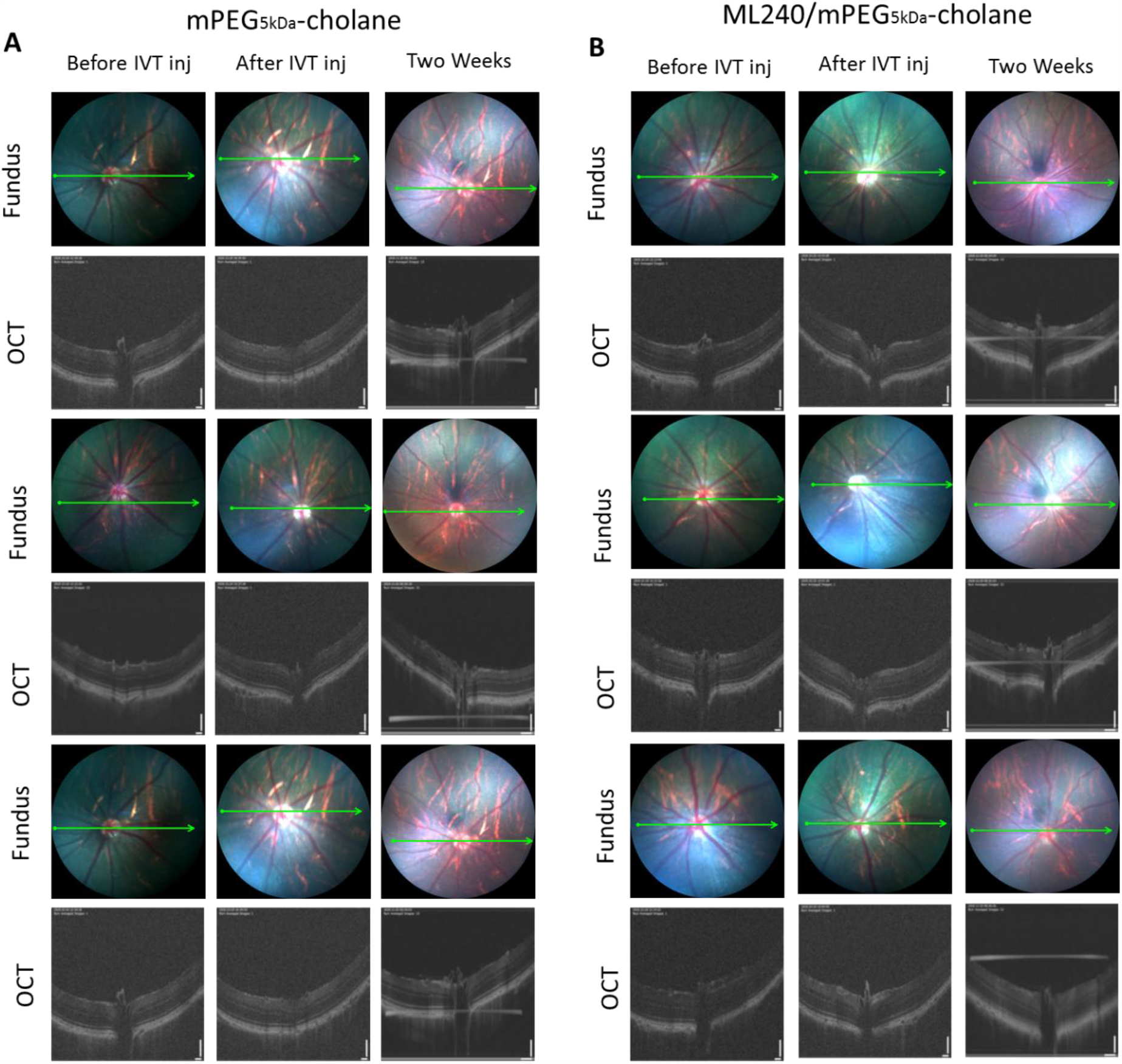
Optical Coherence Tomography (OCT) and fundus images of other replicates of rat vitreous/retina before IVT injection (inj), immediately after IVT injection and following two weeks after IVT injection of (A) drug-free mPEG_5kDa_-cholane nanosystems and (B) 5 µM ML240-loaded mPEG_5kDa_-cholane nanoparticles.

## Notes

### Competing Interest Statement

The authors have declared no competing interest.

